# The potential biofertilization effect of H_2_ is accompanied by a modest impact on the composition of microbial communities in the rhizosphere of common vetch

**DOI:** 10.1101/2025.02.12.637852

**Authors:** Diana P. Dip, Philippe Constant

## Abstract

N_2_-fixing nodules releases molecular hydrogen (H_2_) in the rhizosphere of legumes. The process activates H_2_-oxidizing bacteria (HOB) in soil, leading to multiple effects on biogeochemical processes and a potential biofertilization effect. The legacy effect of the energy potential of H_2_ on soil microbial community structure and the population density of HOB has received little attention. The aim of this study is to evaluate how the legacy effect of HOB, previously activated in soil microcosms exposed to elevated H_2_ concentration (eH_2_), affects biomass production yield of common vetch (*Vicia sativa*) and the abundance of HOB and the composition of microbiome in the rhizosphere. Contrasting soil displaying more than 60% difference in H_2_ oxidation activity were used as growth substrate for vetch. The abundance of HOB was indistinguishable between the treatments and two phylotypes, namely *Aeromicrobium* spp. and *Ramlibacter* spp., were favored by eH_2_ exposure at the activation stage. Their response was transient as no legacy effect was noticed after vetch growth. The root biomass and the root/shoot ratio were reduced in soil conditioned with eH_2_. These results provide new experimental evidence suggesting the biofertilization effect of H_2_ is not universal and requires specific conditions that remain yet to be identified.

## Introduction

With a mixing ratio of 531 ppbv, H_2_ is the second most abundant reducing gas in the atmosphere, after methane (Novelli et al. 1999; Saunois et al. 2019). Besides the atmospheric reservoir, N_2_-fixing nodule of legume plants release H_2_ as an obligatory by-product of N_2_ fixation reaction. In the case of symbiosis involving rhizobia not possessing the NiFe-hydrogenase (*hup*-genotype), the release of H_2_ creates a concentration gradient from approximately 10 000 ppmv at the soil-nodule interface to sub-atmospheric concentration (<0.5 ppmv), 5 cm away from the source (Dong & Layzell 2001; Piché-Choquette et al. 2018). The energy potential of H_2_ is exploited by HOB represented by at least 40 phyla displaying metabolic flexibility (Greening et al. 2016). In carbon limitation conditions, HOB can survive by relying on H_2_ as sole energy source or by a mixotrophic strategy combining H_2_ with organic carbon. It was estimated that 12% of soil bacteria can use H_2_-derived electrons to assimilate CO_2_ (Bay et al. 2021; Xu et al. 2021). Elevated H_2_ concentration in the rhizosphere of legume plants promotes the enrichment of HOB. Incubation of soil samples under modified atmosphere comprising different H_2_ concentrations is a relevant approach to isolate the impact of H_2_ on the composition and activity of soil microbial community. Soil exposure to 250 nmol H_2_ g(soil)^-1^ h^-1^ simulating H_2_ fluxes from N_2_-fixing nodules led to the enrichment of a CO_2_ fixation activity from HOB (Stein et al. 2005). The number of cells affiliated to Cytophaga-Flavobacterium-Bacteroides was also enhanced by H_2_, according to fluorescence *in situ* hybridization analyses. Soil exposure to elevated H_2_ concentrations was also proven efficient to promote PCBs-degrading HOB in soil (Xu et al. 2023). The activation of HOB in soil exposed to H_2_ for a few weeks led variable impacts on the composition of soil microbial communities (Khdhiri et al. 2017; Khdhiri et al. 2018), questioning the incidence of H_2_ on soil microbiota.

The energy potential of H_2_ diffusing in soil exerts effects that are beyond the activity of HOB. That was first exemplified with the higher biomass yield of plants grown in H_2_-fumigated soils when compared to control exposed to ambient air containing trace levels of H_2_. Wheat biomass increased by 48% in soils exposed to a H_2_-enriched atmosphere compared to control soils (Dong et al. 2003). This phenomenon is presented as the H_2_ fertilization effect (Golding & Dong 2010). The exact mechanisms of this fertilization effect remain enigmatic, but some evidence suggests that it involves HOB activated by H_2_. This is supported by indirect evidence relying on the characterization of plant growth promoting activities in some HOB, including *Variovorax* spp. (Maimaiti et al. 2007). The flexibility and metabolic diversity of HOBs could also contribute to the H_2_ fertilization effect through a priming effect on organic matter degradation and nutrient mineralization. There is however an antagonistic effect of HOB activation on the biological sink of atmospheric methane and carbon monoxide, suggesting potential deleterious effects of H_2_ on climate-relevant ecosystem services ((La Favre & Focht 1983; Piché-Choquette & Constant 2019).

This work was conducted to explore if the activation of HOB in soil exposed to an atmosphere containing a high concentration of H_2_ is caused by an increase in their population density. Activation of HOB and alteration of soil microbial community structure would lead biofertilization effect enhancing the growth of *V. sativa*. These hypotheses were tested by setting up a two-stage experimental design. The first stage consisted in conditioning two contrasting HOB populations by exposing a sandy loam soil to either (i) sub-atmospheric (<0.5 ppmv) or (ii) supra-atmospheric (5 000 ppmv) H_2_ concentrations. Soil exposure to H_2_ lasted two months, to favour the activation and enrichment of HOB in soils exposed to high H_2_ concentration. The second stage consisted of growing *V. sativa* using the enriched soils as substrate, without further exposure to modified atmosphere. The plant biomass and microbial communities of the plant rhizosphere were analyzed to examine the legacy effect of H_2_ soil exposure.

## Materials and Methods

### Activation of HOB in soil microcosms

Sandy loam soil was collected in an experimental field trial as described by Agoussar et al. (2021). Soil was homogenized (2 mm sieve), the water content was adjusted at 30% water holding capacity and a defined amount (300 g) was transferred into 500 mL (nominal) Wheaton borosilicate bottles, occupying approximately half of the total volume. Bottles were sealed with a gas tight rubber septum cap. Two different H_2_ treatments were applied: elevated H_2_ concentration (eH_2_) and sub-atmospheric H_2_ concentration (control). Soil exposure to eH_2_ was achieved by injecting a defined volume of certified 99,999% compressed H_2_ (Praxair Distribution Inc., PA, USA) in bottles to reach 5000 ppmv in the static headspace. Injections were conducted every 2-3 days after 30 minutes of equilibration of the headspace with the ambient atmosphere. For the control treatments, bottles were left open for 30 min to equilibrate the headspace with atmospheric H_2_ (approximatively 0.5 ppmv). Replacement of the static headspace was done every 2-3 days. Microcosms were incubated for two months in dark conditions, at room temperature to activate HOB (de la Porte et al. 2024; Piché-Choquette et al. 2016). Each experimental unit was represented by 20 repetitions (total of microcosms = 40). Soil subsamples 0.25 g) were collected after the incubation for DNA extraction and the balance was transferred in pot for vetch cultivation.

### Hydrogen oxidation rate

H_2_ oxidation rate was monitored sporadically, after seven incubation days. Measurements were performed on a random selection of three soil microcosms representative of each H_2_ exposure treatment. A chromatographic assay was utilized to measure the first-order oxidation rate under 5000 ppmv H_2_ concentration (de la Porte et al. 2024). Calibrations were conducted using a certified H_2_ gas mixture (10 000 ppmv H_2_, balance air). Linear regression was integrated from three dilutions of the certified gas mixture, including 0.1% (1000 ppmv), 0.5% (5000 ppmv) and 1% (10 000 ppmv) H_2_.

### *Vicia sativa* pot experiment

Soil microcosms representative of each H_2_ exposure were randomly selected after the activation stage (10 soil samples x 2 treatments = 20 pots in total). Soil samples (approximately 300 g) were mixed with vermiculite (5:1, soil:vermiculite) and transferred in 1 L pots. *V. sativa* L. seeds (EBENA BRAND-FF 2023-0261, Meuneries Mondou) were surfaced disinfected and allowed to germinate in petri dishes containing 1% Water-Agar solution at 25°C, 50% with a photoperiod of 16h/8h light/darkness, respectively. Seedlings were transferred into the pots and randomly placed in a growth room, under full spectrum Led lights for two months. Pots were regularly watered with tap water. The biomass of *V. sativa* was harvested for the determination of fresh and dry weight of aerial and roots biomass. The rhizosphere soil was collected for subsequent DNA extraction.

### PCR amplicon sequencing of bacterial 16S rRNA gene

Total DNA was extracted from soil microcosms after the activation stage and from rhizosphere soil after plant harvest with the PowerLyzer PowerSoil kit (Qiagen), following the instructions from the manufacturer. PCR amplicon sequencing of bacterial 16S rRNA gene was performed with the primers 515F and 806R, targeting the V4 hypervariable region of 16S rRNA gene (Caporaso et al. 2011). Raw sequence reads were deposited in the Sequence Read Archive of the National Center for Biotechnology Information under BioProject PRJNA1200744. Procedure for library preparation was followed as described by Saavedra-Lavoie et al. (2020). The quality control and sequencing were done at the *Centre d’expertise et de service Génome Québec* (Montreal, Québec, Canada), with the Illumina MiSeq PE-250 platform. Downstream analyses of raw sequences were performed using the cutadapt, for primer removal, and Dada2 pipeline, for quality control, amplicon sequence variant (ASV) table and taxonomic assignation, with the RStudio software (Callahan et al. 2016; Martin 2011). The taxonomic assignation was based on SILVA v128 database (Quast et al. 2012). ASV representing less than 0.005% relative abundance were not considered for downstream analysis. Species richness, Shannon and Simpsons index were obtained with the package *iNEXT* v3.0.1 (Chao et al. 2014; Hsieh et al. 2016).

### Quantification of *hhyL* gene abundance

The abundance of *hhyL* gene, encoding for the large subunit of NiFe-hydrogenases group 5/1h, was determined in soil samples collected during the activation stage. The droplet digital PCR (ddPCR) assay described by Baril et al. (2022) was utilized. Randomly selected DNA from each treatment (control and eH_2_) were used for the ddPCR assays. Negative controls were tested by using with DNA-free sterile water. Threshold was set manually to include rain in the positive fraction (considering only those with up until 12% error), and samples with more than 10 000 droplets were selected for further analyses. Copy number concentrations were converted to copy per gram of dry soil.

### Statistical analyses

Statistical analyses were performed using the software Rstudio v4.2.3 (Team 2013). Graphics were generated with the package *ggplot2* v3.5.2. ASVs aggregated at the genus level were subjected to ANCOM-BC analyses to assess the response of bacteria to the activation phase, using the *microbiome* v1.20 and *ANCOMBC* v2.0.3 packages (Lahti & Shetty 2018; Lin & Peddada 2020). Comparison of biomass DW between treatments and alpha diversity of microbial communities, was examined with one-way ANOVA using de stats package v4.2. A t-test was computed to compare the absolute abundance of *hhyL* genes between the two H_2_ exposure treatments. Raw datasets utilized for statistical analyses are provided as supplementary material (Supplementary Table S1).

## Results

### HOB activation stage

The activation of HOB conducted in this study was prolonged to 10 weeks to verify whether extension of H_2_ exposure can support short-term proliferation of HOB through mixotrophy or chemosynthesis. H_2_ oxidation rate was measured from three independent replicates of each treatment, from seven days to 71 days exposure to modified atmosphere. *Figure 1* shows, as expected, that the H_2_ oxidation rate measured in control treatment was relatively stable, at 29±3 nmol g_(soil-dw)_^-1^ h^-1^ on average, without significant temporal pattern. Activation of HOB in microcosms exposed to elevated H_2_ concentration was efficient with H_2_ oxidation rate varying from 47±8 to 80±11nmol g_(soil-dw)_^-1^ h^-1^ after 30 day of exposure to 63±5 nmol g_(soil-dw)_^-1^ h^-1^ at the end of the incubation period. The plateau of H_2_ oxidation rate indicates the energy potential of H_2_ and nutrient content in soil were not sufficient to support proliferation of HOB. The absence of proliferation is supported by indistinguishable *hhyL* gene copy numbers among control with 2.2±1.7 ×10^6^ copies per g, and H_2_ treatments with 2.9± 2.2 ×10^6^ copies per g (t-test, p-value = 0.56).

**Figure 1.**
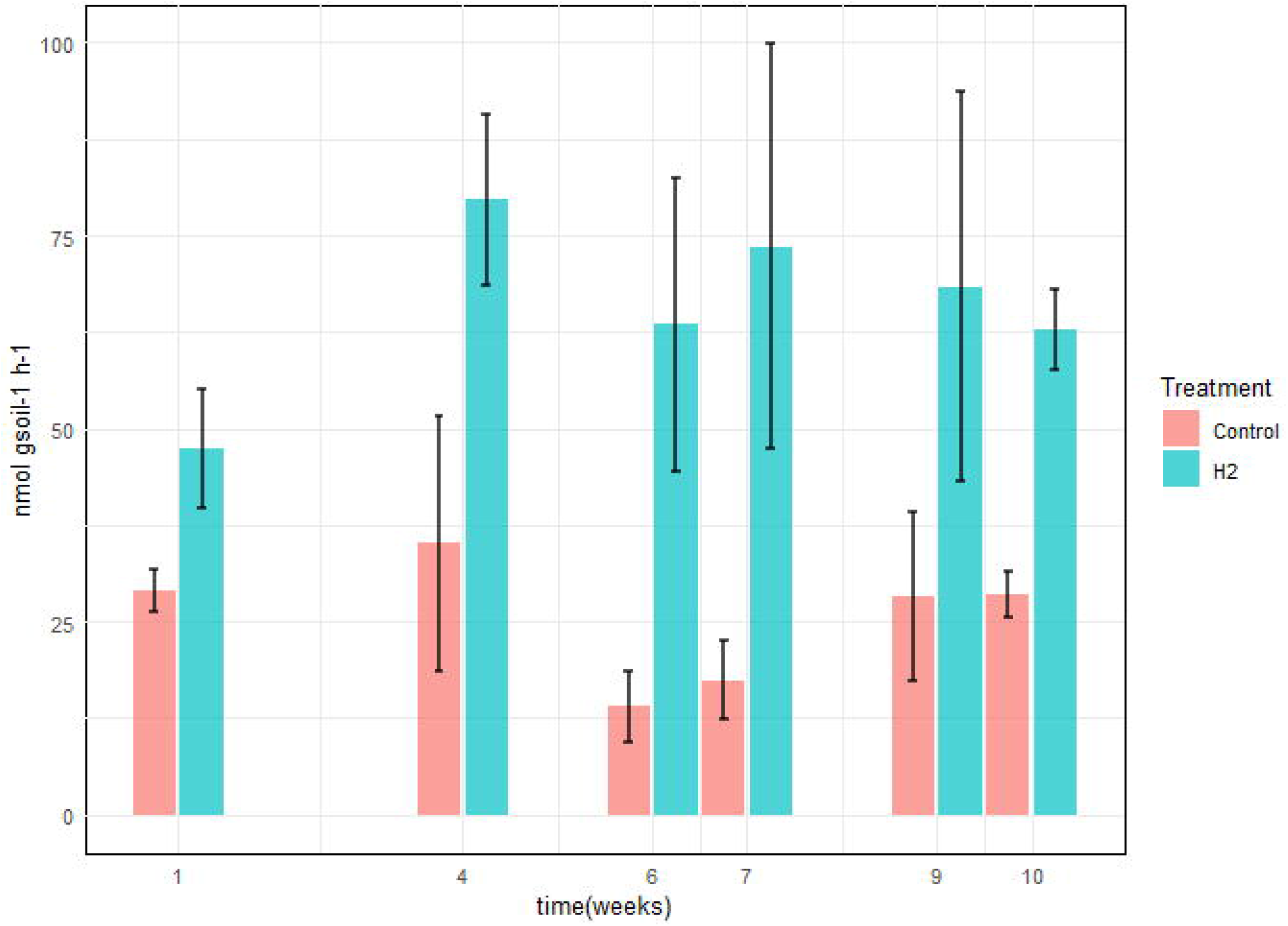
H_2_ oxidation rates in soil microcosms exposed to elevated (H_2_) or sub-atmospheric (Control) H_2_ concentrations. Each measurements routines were conducted on three randomly selected replicates, during the activation stage of the experiments.

### Soil microbial communities

Most bacterial 16S rRNA gene sequences were affiliated to the phyla Actinobacteria (28%), Proteobacteria (27%), Thaumarchaeota (16%) and Acidobacteria (10%). The impact of H_2_ exposure during the activation stage on soil bacterial communities was negligible, as variations of alpha diversity were not significantly influenced by the H_2_ treatment (Table S1). Aggregation of the ASV at the genus level led to the identification of three genera displaying contrasting differential abundances among both H_2_ exposures. Two groups were favored by H_2_ exposure, namely *Aeromicrobium* spp. (LFC = 1.02, p-value = 0.002) from the Actinobacteria phylum and *Ramlibacter* spp. (LFC = 0.74, p-value = 0,00005) from the Proteobacteria phylum (*Fig. 2*). A single group of ASV aggregated to the genus *Pseudoduganella* spp. appeared more favored in the control treatment than in the H_2_ treatment but the difference was not significant (LFC = - 1.25, p-value = 0.08). No legacy effect of H_2_ exposure was observed on soil microbial communities in the rhizosphere of vetch as neither the alpha diversity nor the distribution of ASV aggregated at the genus level was influenced by H_2_ exposure treatment.

**Figure 2.**
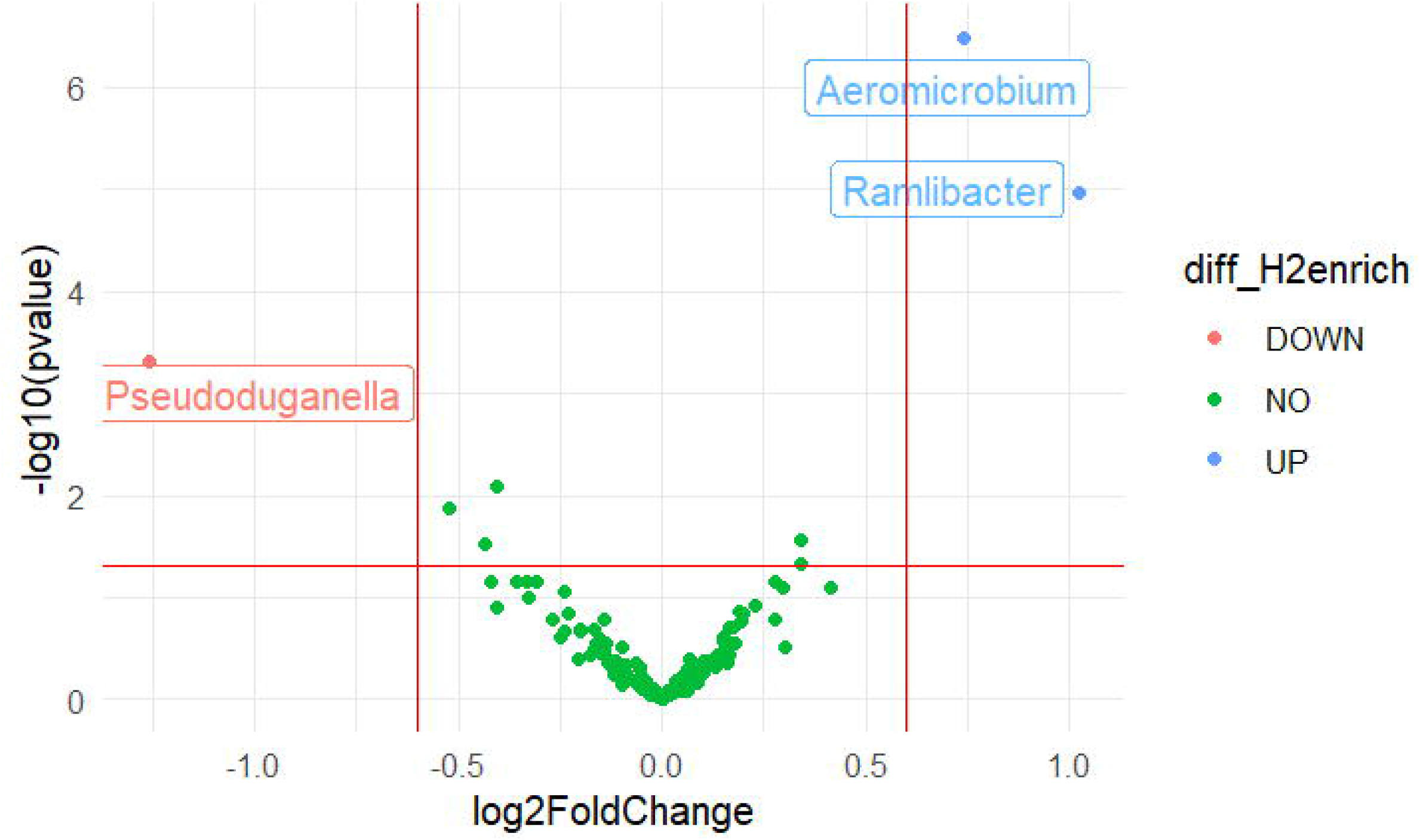
Volcano plot representing the ANCOMBC analysis for the activation stage of microbial communities. log_2_ fold change (eH_2_/Control) of ASV aggregated at the genus level in response to H_2_ exposure at the activation stage of the experiments. Blue dots represent genera favored by eH_2_ soil exposure, and the red dot represents the single genus favored in the Control treatment. Green dots show non-responsive bacterial genera to H_2_ exposure (ANCOMBC, n=10).

### The legacy of HOB activation of vetch

The potential fertilization effect of H_2_ was examined on aerial and below-ground biomass of vetch (*Fig. 3*). The aerial biomass of *V. sativa* was not influenced by H_2_ treatment whereas roots biomass was reduced in plants growing in soil conditioned under eH_2_. The root/shoot ratio, usually used as an observation of the plant response to the environment, was reduced in eH_2_ exposure treatment (p-value = 0.028).

**Figure 3.**
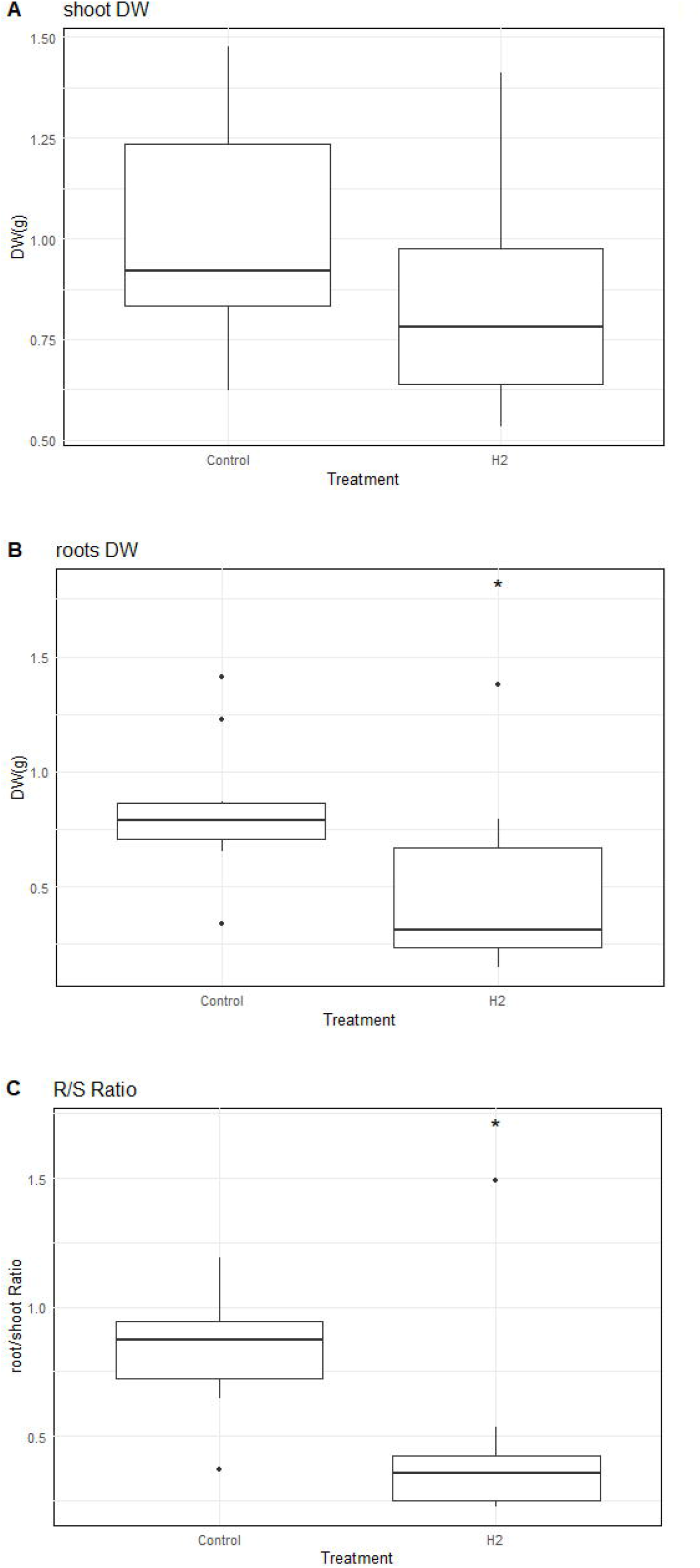
Differences in dry weight of *V. sativa* after harvest in soil conditioned under elevated (H_2_) or sub-atmospheric (Control) H_2_ concentrations. The dry weight of (A) aerial, (B) root, and (C) root/shoot ratio is represented (n=10).

## Discussion

The production of H_2_ in N_2_-fixing nodule comprising rhizobia not possessing [NiFe]-hydrogenase creates steep H_2_ concentration gradient in the rhizosphere of legume plants. These concentrations gradients expand at long distances, up to several centimeters away from nodules. In addition to redefine the spatial boundaries of the rhizosphere effect, these gradients are expected to play a role in modifying microbial community structure and function (de La Porte et al. 2020). This notion was first supported by the enrichment of HOB activity in soil surrounding N_2_-fixing nodules in pioneering investigations performed in pot experiments. Subsequent efforts were invested in artificial soil microcosms exposed to elevated H_2_ concentrations. H_2_ exposure enhanced net CO_2_ fixation in soil, which is due to the facultative chemolithoautotrophic life strategy of numerous HOB in soil (Stein et al. 2005). The application of terminal restriction fragment length polymorphism (T-RFLP) led to the conclusion that only a few members were favored by H_2_ exposure (Osborne et al. 2010). This observation was contrasted in subsequent investigations applying high-throughput PCR amplicon and metagenomic sequencing techniques coupled with loose differential abundance analyses, suggesting H_2_ exposure induced significant alteration of microbial communities, mainly on the rare biosphere (Khdhiri et al. 2017; Piché-Choquette et al. 2016). Revisiting these experiments with more stringent statistical tools tailored to sparse microbiome data here, and in a recent work (de la Porte et al. 2024), brings us back to conclusions supported by earlier T-RFLP techniques, namely H_2_ exerts neglecting impact on soil microbial community. The exact mechanisms responsible for biogeochemical processes variations upon soil exposure to H_2_ are therefore decoupled from HOB proliferation and compositional change of microbial communities.

Only two ASV groups were positively influenced by the H_2_ exposure during the activation stage (*Aeromicrobium* and *Ramlibacter*). Pumphrey et al. (2011), reported the enrichment of labelled DNA from a phylotype affiliated to *Aeromicrobium ginsengisoli* from hairy vetch rhizospheric soil incubated with 100 ppm of H_2_. There is therefore evidence for chemolithoautotrophic metabolism in *Aeromicrobium* spp. but demonstration of the activity in isolates has not been reported. The negligible alteration of microbial communities by H_2_ is paralleled with no evidence supporting the proliferation of HOB in soil exposed to H_2_. The stabilization of the H_2_ oxidation activity in soil exposed to H_2_ was suggested in previous microcosms experiment where the response of high-affinity H_2_ oxidation activity plateaued after one-week H_2_ exposure period (Piché-Choquette et al. 2016). Extended incubation under a headspace enriched with H_2_ was insufficient to induce a secondary enrichment after the enrichment plateau. The level of CO_2_ in the static headspace of soil microcosms during the activation stage as well as organic carbon and other nutrients were likely insufficient to support the proliferation of HOB.

Another aspect of H_2_ exposure in agroecosystem is the fertilization effect of this gas. This was first reported in pot experiments where soil substrate was fumigated with H_2_, prior to plant seedling (Dong et al. 2003). The plant biomass increased by 15–48% in fumigated soil compared to controls. The fertilization effect was attributed to HOB displaying plant growth promotion effect (Maimaiti et al. 2007), such as 1-aminocyclopropane-1-carboxylate (ACC) deaminase activity reducing the accumulation of ethylene playing a role of stress phytohormone (Yang & Hoffman 1984). This is thus indirect evidence suggesting that recruitment of symbiotic N_2_-fixing rhizobia releasing H_2_ (Hup^-^) is an evolutionary advantage for legume plants. Rhizobia possessing [NiFe]-hydrogenase (Hup^+^) to recycle H_2_ produced by nitrogenase exist, but they appeared less represented by their Hup^-^ counterparts in nature (Annan et al. 2012). Consideration of the fertilization effect of H_2_ and the selection of Hup-symbioses in nature led to the suggestion that H_2_ is a missing link explaining the multiple benefits of crop rotation that was not reachable by nitrogen fertilization alone (Golding & Dong 2010).

Benefits of H_2_ on plants are likely multifactorial. The fertilization effect was not observed for wheat grown in soil subjected to fumigation by H_2_, CO or CH_4_ (de la Porte et al. 2024). The aerial biomass of vetch in the present study was not enhanced in soil conditioned with H_2_ exposure. The reduction in root biomass, as well as in root/shoot ratio, is thought to be related to plant growth promotion activity of HOB, including ACC deaminase reducing root proliferation for nutrient prospection. If this mechanism holds true, it was realized in the absence of evident HOB proliferation. These results must be considered in trials integrating irrigation treatments with H_2_-satured water enhancing stress resistance in plants. The antioxidant potential of H_2_, microbial activation of immune system in plants and HOB are potential contributors to plant benefits (Wang et al. 2024). A more holistic analysis examining both plant and microorganisms interactions in soil exposed to different H_2_ level or in contrasting Hup+/Hup-symbioses are recommended to disentangle the contribution of microorganisms and plant metabolism in explaining the benefit of H_2_.

## Conclusions

In conclusion, soil exposure to eH_2_ was not sufficient to support the growth of HOB as we hypothesized. H_2_ was likely sufficient to support persistence energy requirement of HOB, with a rewiring of energy metabolism toward inorganic energy source and a repression of carbon metabolism (Greening et al. 2015; Liot & Constant 2016). The supply of certain carbon sources along with H_2_ could promote growth and increase activity of HOB in soil (Baril & Constant (2023). Examination of potential synergetic effect of H_2_ with nutrients will decipher the limiting factors for HOB proliferation and activation in soil.

## Acknowledgments

This work was supported by a Discovery grant (RGPIN-2024-06451) from the Natural Sciences and Engineering Research Council of Canada (NSERC) to P.C.

